# Sex differences in neurotransmitter levels in different brain regions after acute and chronic morphine treatment in mice

**DOI:** 10.1101/2023.01.16.524193

**Authors:** Florian Gabel, Volodya Hovhannisyan, Abdel-Karim Berkati, Virginie Andry, Yannick Goumon

## Abstract

**Background and Purpose:** Pain management is a major health burden. Pain results from the integration of the nociceptive message and neuronal communication relying on neurotransmitters such as glutamate, γ-aminobutyric acid (GABA), dopamine, noradrenaline and serotonin in brain regions including the periaqueductal gray (PAG), the nucleus accumbens (Nac), the caudate-putamen (Cpu) and the amygdala. Morphine remains the gold standard painkiller for severe pain *via* the activation of the mu opioid receptors. However, among side effects, morphine chronic treatment lead to antinociceptive tolerance. As antinociceptive tolerance might be linked to neurotransmission dysregulation, we have compared various neurotransmitter concentrations in acute *vs* chronic morphine conditions in the amygdala, PAG, Cpu, and the Nac of male and female mice.

**Experimental approach:** Sex differences in morphine antinociception and tolerance were assessed using the tail-immersion test. The behavioural effects of acute and chronic morphine treatments, as well as sex differences in the levels of dopamine, serotonin, noradrenaline, glutamate and GABA in the amygdala, PAG, Cpu, and the Nac were determined by an absolute quantification LC-MS/MS approach using the isotopic dilution method.

**Key results:** This study indicates, as previously reported, that female mice are less sensitive to morphine and develop morphine antinociceptive tolerance earlier than males (ED50 of 5.5±0.24 days *vs* 1.54±0.11 days, respectively). However, the rate at which the tolerance developed did not differ between both sex. We have found major differences in dopamine, serotonin, noradrenaline, glutamate and GABA levels between female and male mice in the amygdala, PAG, Cpu, and the Nac. Finally, no major effect of anti-nociceptive tolerance induced by chronic morphine was observed compared to acute administration of morphine.

**Conclusion:** Neurotransmitter differences are attributable mainly to sex differences in pain-related CNS regions. However, the impacts of morphine anti-nociceptive tolerance on dopamine, serotonin, noradrenaline, glutamate and GABA contents appeared to be limited.

## INTRODUCTION

Nowadays, pain management represents one of the most important health challenges as chronic pain has become one of the first cause of disability in the world [1]. In humans, pain is a complex sensory and emotional experience tightly influenced by biological, psychological and social factors [2]. From a physiological point of view, pain arises from the integration of the nociceptive message that occurs in different regions of the central nervous system (CNS) such as the periaqueductal gray (PAG), the basal ganglia (Nucleus accumbens, Nac; caudate-putamen, Cpu), the amygdala and cortical regions including the prefrontal cortex and the anterior cingular cortex (for review: [3–6]). In these structures, the transmission of the nociceptive signal relies on neurotransmitters including dopamine, serotonin, noradrenaline, *γ*-aminobutyric acid (GABA) and glutamate which play important roles in pain processing (for review: [7–9]).

In the clinic, a range of treatments can be used according to pain severity. Morphine remains the gold-standard painkiller to treat moderate to severe pain despite its numerous side effects [10]. Morphine antinociception is mostly the result of its binding to mu opioid receptors (MORs) located on neurons of the CNS. These receptors are found throughout the CNS, but those expressed in regions related to pain such as the PAG and the amygdala are especially crucial for antinociception [11–13]. In these structures, morphine is involved in the modulation of GABA and glutamate transmission ultimately leading to antinociception (For review, see [6, 14, 15]). Interestingly, other neurotransmission systems have been shown to modulate morphine antinociceptive effects. For instance, serotonin reuptake inhibitors and dopaminergic signaling within the PAG, enhance morphine antinociception [16, 17].

Among morphine side effects, antinociceptive tolerance corresponds to the loss of morphine effect after chronic administration. Consequently, dose escalation is needed to achieve sufficient antinociception despite a higher risk of addiction and overdose [10, 18]. Chronic morphine administration has been shown to modulate neurotransmitter systems in the brain [19, 20]. For instance, MOR signaling after chronic administration of morphine has been described to induce an increase of dopaminergic transmission participating in the development of addiction [21, 22], while modulation of serotonin neurotransmission is involved in both morphine dependence (occurring after a chronic treatment) and withdrawal [23]. In addition, the expression of numerous receptors involved in neurotransmission is also regulated by chronic morphine administration. Indeed, the expression of the 5-hydroxytryptamine receptor 2C (5-HT_2C_) in the ventral tegmental area (VTA), the locus coeruleus and the Nac in mice [24], β-adrenergic receptors expression within the rat parietal cortex [25], NMDA receptors expression in the rat amygdala [26], as well as GABA_B_ receptor subunits 1C and 1D expression in the locus coeruleus and VTA of mice and rats are upregulated during a chronic treatment of morphine [27].

Interestingly, neurotransmitters including serotonin, noradrenaline [28], GABA [29] as well as glutamate [30], are involved in the dopaminergic system and indirectly modulate various aspects of addiction. Their modulation by morphine might, therefore, alter the dopaminergic transmission and addiction.

We have recently shown that morphine antinociception and tolerance, as well as its metabolism in the brain, are influenced by sex [31]. In the brain, morphine is metabolized into its predominant metabolite M3G which is believed to oppose morphine effects (*i.e*., produces hyperalgesia) [32]. Although M3G mechanism of action remains unclear, several studies have suggested an alteration of neurotransmission as a consequence of M3G effects (for review, see Gabel 2022). In addition, even though we have shown that morphine metabolism does not seem to be altered after chronic treatment, the repeated presence of morphine and/or M3G in the brain might modulate neurotransmitter production in a sex-specific manner. To our knowledge, simultaneous comparison of various neurotransmitter concentrations in acute *vs* chronic morphine conditions in male and female mice has never been performed. Hence, the present article investigates differences in neurotransmitter levels in the amygdala, PAG, Cpu, and the Nac after an acute or chronic morphine treatment leading to antinociceptive tolerance in male and female mice.

## MATERIALS AND METHODS

### Animals

Experiments were performed with ten weeks-old male and female C57BL/6J mice (26±4 g and 20±4 g, respectively; JAX:006362; Charles River, L’Arbresle, France). Male and female mice were group-housed at 5 per cage, with a 12 h light-dark cycle, at a temperature of 23°C±2°C and provided with food and water *ad libitum*. All procedures were performed following European directives (2010/63/EU) and were approved by the regional ethics committee (CREMEAS) and the French Ministry of Agriculture (license No. APAFIS# 23671-2020010713353847v5 and APAFIS#16719-2018091211572566v8 to Y.G.). The ARRIVE Guidelines have been followed for reporting experiments involving animals [33].

### Behavioural assessment of morphine antinociceptive effect

The antinociceptive effect of morphine was measured with the tail immersion test (TIT) as described before [31]. Briefly, mice were first accustomed to the experimental conditions. Then, they were tested by measuring the latency of the tail withdrawal when 2/3 of the tail was immersed in a constant-temperature water bath heated at 47°C. In the absence of a response, the cut-off was set at 25 s to avoid tissue damage. Results are expressed as % maximal possible effect (MPE) according to the following formula:

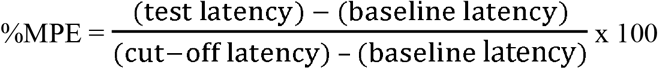

### Induction of morphine antinociceptive tolerance

Mice were injected i.p. with 10 mg/kg of morphine (2 mg/ml; volume of 5 μl/g of mouse; Francopia, Paris, France) every morning (light phase at 10 AM) for nine consecutive days. Morphine was dissolved in saline solution (NaCl 0.9%, w/v) and control mice were injected i.p. with an equal volume of saline solution. Mice were tested before and 30 min after each injection. On day 10, all mice received an i.p. injection of 10 mg/kg of morphine before the final procedure.

### Brain regions sampling

On day 10, mice were sacrificed by cervical dislocation 30 min following the injection of morphine and the brain was collected and placed on an ice-cold mouse brain matrix. Razor blades were used to cut the brain into 1 mm thick slices. A puncher of 1 mm diameter was used to sample the PAG and the Cpu while a puncher of 0.5 mm diameter was used for the amygdala and the Nac. Brain regions were directly transferred into micro-tubes and stored at −80°C.

### Sample preparation

Tissues were homogenized in 200μL of 0.1 mM ascorbic acid, sonicated 2 x 5 sec at 100W and then centrifuged (20,000g, 30 min, 4°C). Supernatants were recovered and protein concentrations were determined with the Bradford method (Protein Assay kit, Bio-Rad, Marnes-la-Coquette, France). Then, samples were derived with the AccQtag Ultra derivatization kit (Waters, Saint-Quentin-en-Yvelines, France). Briefly, 10μl of each sample was mixed with 30 μl of borate buffer and 10 μl of internal standards (IS; 80 pmol of D4-dopamine, 80 pmol of D4-serotonin, 80 pmol of D6-noradrenaline, 100 pmol of D6-GABA and 360 pmol of D5-glutamate; Sigma Aldrich, Saint Quentin Fallavier, France). Then, 10μl of AccQtag reagent was added and the mixture was incubated for 10 min at 55°C under shaking (500 rpm). After the incubation, 200ul of acetonitrile 100% was added to the samples and samples were shaken, centrifuged (20,000g, 4°C, 5 min) and dried under vacuum. Samples were resuspended in 20 μl of H2O/0.1% formic acid (v/v) and 5μl were analysed by liquid chromatography-tandem mass spectrometry (LC-MS/MS).

### LC-MS/MS instrumentation and analytical conditions

Analyses were performed with a Dionex Ultimate 3000 HPLC system (Thermo Electron, Villebon-Sur-Yvette, France) coupled with a triple quadrupole Endura mass spectrometer (Thermo Electron). Xcalibur v4.0 software (RRID: SCR_014593; Thermo Electron was used to control the system. Samples were loaded onto a ZORBAX SB-C18 column (150 x 1 mm, 3.5 μm, flow of 90 μl/min; Agilent, Les Ulis, France) heated at 40°C. LC and MS/MS conditions used are detailed in **Table S1**.

Identification of the compounds was based on precursor ions, selective fragment ions and retention times obtained for the heavy counterpart present in the IS. Selection of the monitored transitions and optimisation of collision energy and RF Lens parameters were determined manually (for details, see **Table S1**). Qualification and quantification were performed using the multiple reaction monitoring mode (MRM) according to the isotopic dilution method [34].

### Data and Statistical Analysis

Statistical analysis was performed using GraphPad Prism 6 Software (RRID: SCR_002798). To compare the groups, two-way ANOVA was used and followed by Tukey’s multiple comparisons test only if F was significant and there was no variance homogeneity. Normality and variance homogeneity were checked with the D’Agostino & Pearson omnibus normality test and Levene’s test, respectively. If assumptions were violated, a non-parametric approach, the Aligned-Rank Transform (ART) ANOVA, was used [35].

For the behavioural experiment, non-linear regression with a 4-parameters logistic equation was used to define the average day at which half of the MPE remains and the Hill slope coefficient in both males and females. The means of each parameter were compared using an unpaired t-test after that normality and variance homogeneity were checked. Results are presented as mean values ± standard error of the mean (SEM).

## RESULTS

### induction of morphine antinociceptive tolerance

To investigate the impact of chronic morphine treatment, we first used the tail-immersion test (TIT) to assess the development of morphine antinociceptive tolerance in male and female mice (see protocol **Figure 1a**). Statistical details are presented in **Table S2**.

**Figure 1.**
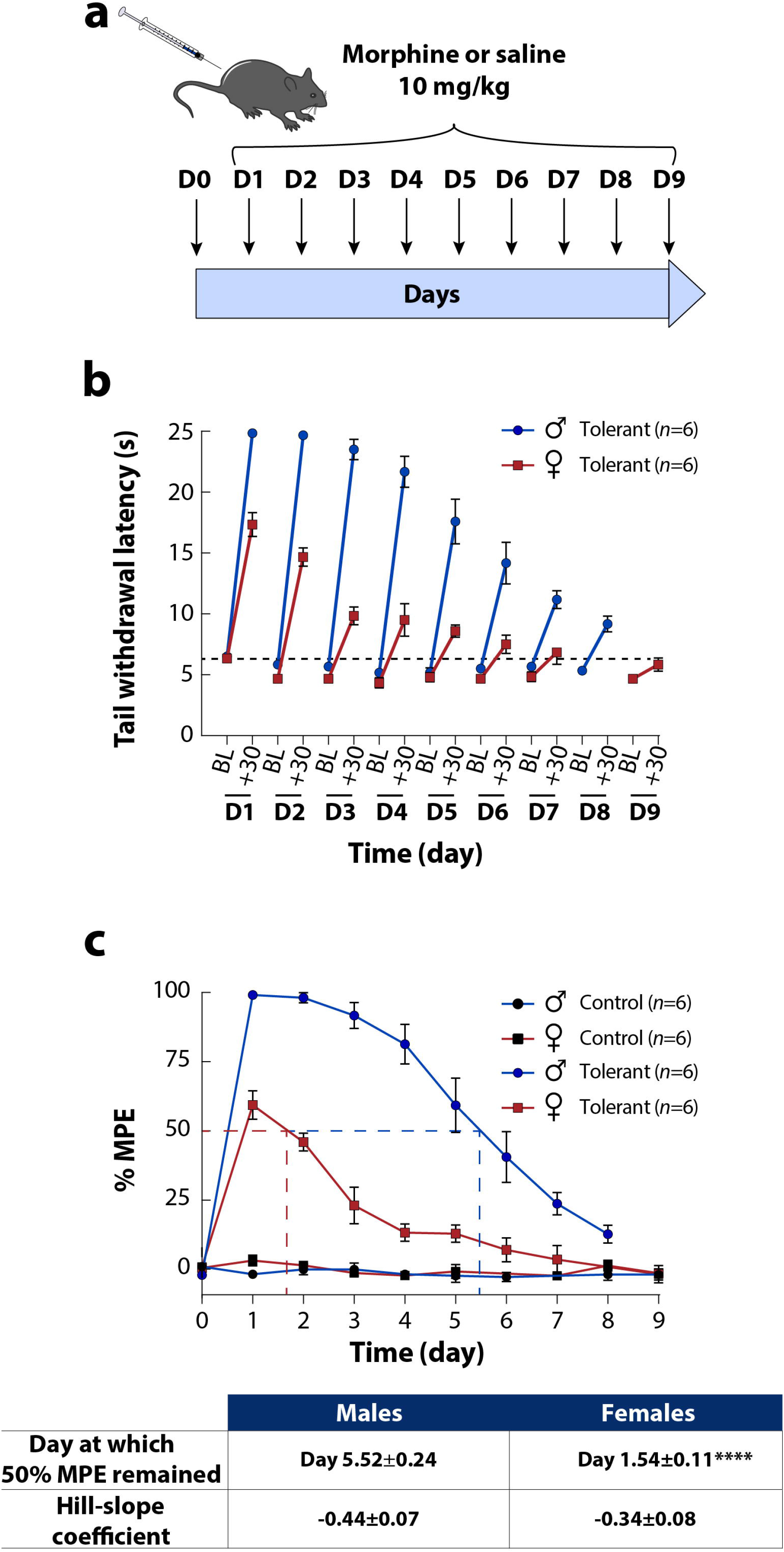
Antinociceptive effect of morphine and development of morphine antinociceptive tolerance in male and female mice. (**a**) Protocol of induction of the antinociceptive tolerance to morphine. (**b**) Development of morphine antinociceptive tolerance throughout the chronic treatment (i.p. 10 mg/kg, from day 1 to 9). Tail withdrawal latency was observed 30 min after morphine or saline injection for 9 successive days. Data are expressed as mean ±SEM; *n* are indicated in the figure. (**c**) Development of morphine antinociceptive tolerance throughout the chronic treatment (i.p. 10 mg/kg, from day 1 to 9). Control groups correspond to mice treated only with saline vehicle. Antinociception is expressed as %MPE observed before and 30 min after morphine or saline injections for 9 successive days. Data are expressed as mean ±SEM; *n* are indicated in the figure. ED_50_ were extracted from each fitting, and fits were compared with the extra sum-of-the-square F test. Unpaired t-test with Welch’s correction was used to compare the values for the day at which half of the MPE remains and the hill-slope coefficient values. ****, *P*<0.0001. Males are represented as blue/black circle dots and a blue line, whereas females are represented as red/black square dots and a red line.

The tail-withdrawal latencies measured throughout the protocol decreased over the course of the treatment in both males and females. There were no sex differences in baseline nociceptive thresholds of the mice **(Figure 1b)**. After the first injection, the MPE of morphine decreased gradually over the time course in both male and female mice, consistent with the induction of antinociceptive tolerance (**Figure 1c**). Interestingly, males reached the upper cut-off of the tail-immersion test (25s) on the first three days of the treatment, whereas female mice did not reach the cut-off even on day 1. In addition, females became tolerant to morphine antinociceptive effects earlier than males. Hence, significant differences were observed between males and females on the day at which only 50% of MPE remains (**Figure 1c**). However, no difference was observed in the Hill slope coefficients, indicating that the rate at which the tolerance appeared seems to be identical in males and females (**Figure 1c**). One should yet note that sex differences in Hill slope coefficients may have been masked by the cutoff used during the experiment (i.e., 25 s).

These results are consistent with our previous observations in mice and suggest that females are less sensitive to morphine antinociception and become tolerant to morphine earlier than males [31].

### Quantification of dopamine, serotonin, noradrenaline, GABA and glutamate in brain regions

Neurotransmitter levels in the brain are known to be influenced by sex [36–38], as well as by chronic treatment with morphine [39, 40]. However, to our knowledge, the interaction between sex and chronic morphine treatment was never assessed before. We have thus investigated the consequences of acute and chronic morphine treatment on neurotransmitter levels in different brain regions in male and female mice. Four brain regions were analysed: the amygdala, the PAG, the Cpu and the Nac. Dopamine, serotonin, noradrenaline, GABA and glutamate levels were quantified by LC-MS/MS. Statistical details are presented in **Table S2**.

For the reader’s convenience, the control group (CT) refers to the acute treatment with morphine, while the tolerant group (TOL) designates the chronic treatment with morphine leading to antinociceptive tolerance. In addition, to better visualize the overall sex differences observed in all the structures, the normalized quantities (according to male levels for each neurotransmitter) of dopamine, serotonin, noradrenaline, GABA, and glutamate, as well as the normalized GABA/glutamate ratio are illustrated in CT (**Figure 2g, 3g, 4f, 5f**) and TOL mice (**Figure 2h, 3h, 4g, 5g**).

**Figure 2.**
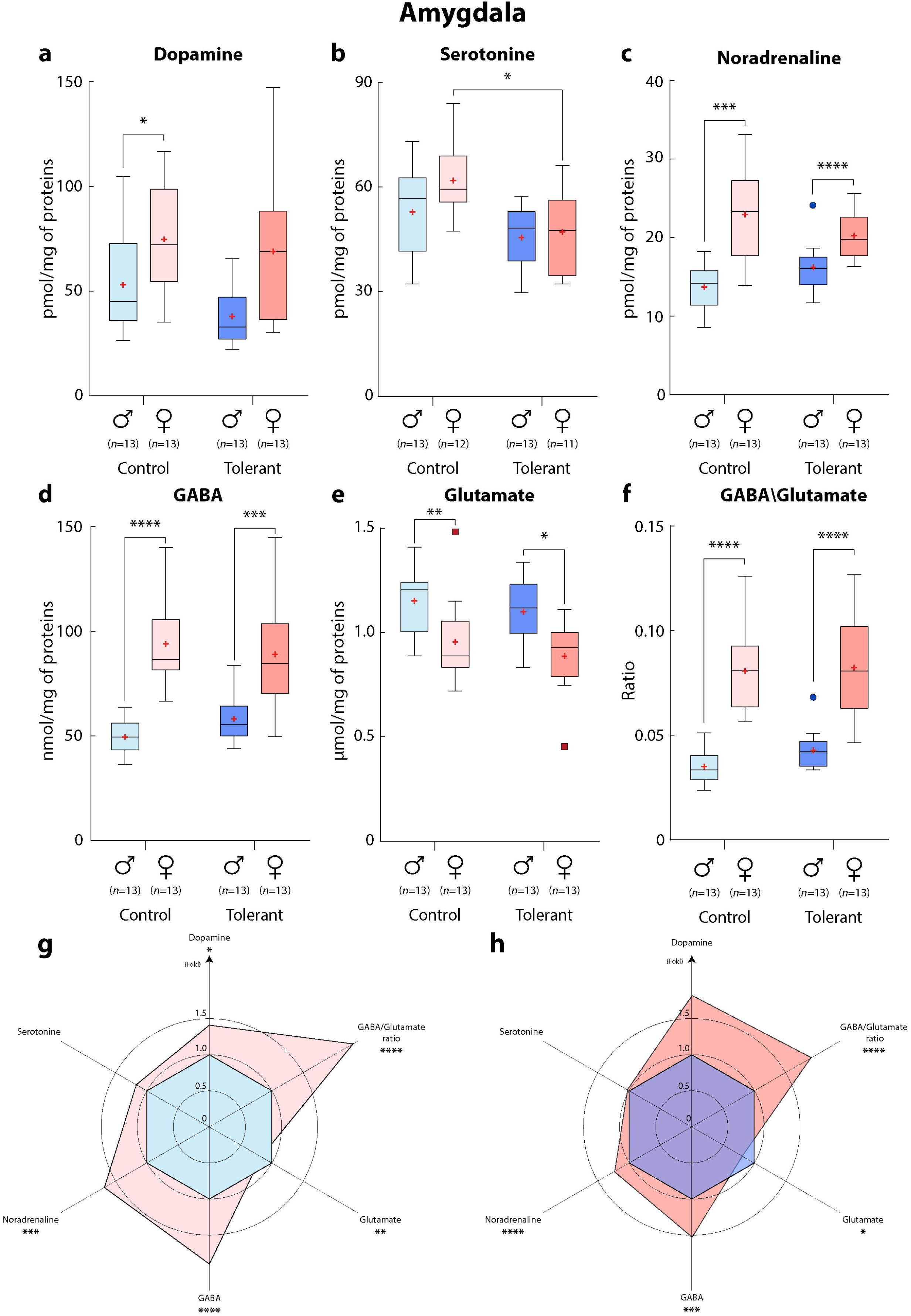
Concentrations of neurotransmitters in the amygdala of male and female mice treated with acute or chronic morphine. (**a**) dopamine, (**b**) serotonin, (**c**) noradrenaline, (**d**) GABA, (**e**) glutamate, (**f**) GABA/glutamate ratios. Normalized quantities (according to male levels for each neurotransmitter) of dopamine, serotonin, noradrenaline, GABA, and glutamate, and GABA/glutamate ratio are presented in (**g**) CT mice and (**h**) TOL. The control group (CT) refers to the acute treatment with morphine, while the tolerant group (TOL) designates the chronic treatment leading to antinociceptive tolerance. Data are expressed as means ± SEM, *n* are indicated in the figure. Two-way ANOVA was applied and followed by Tukey’s multiple comparisons test only if F was significant and there was no variance homogeneity. Tukey’s multiple comparisons results are reported as *, *P*< 0.05; **, *P*<0.01; ***, *P*<0.001; ****, *P*<0.0001.

**Figure 3.**
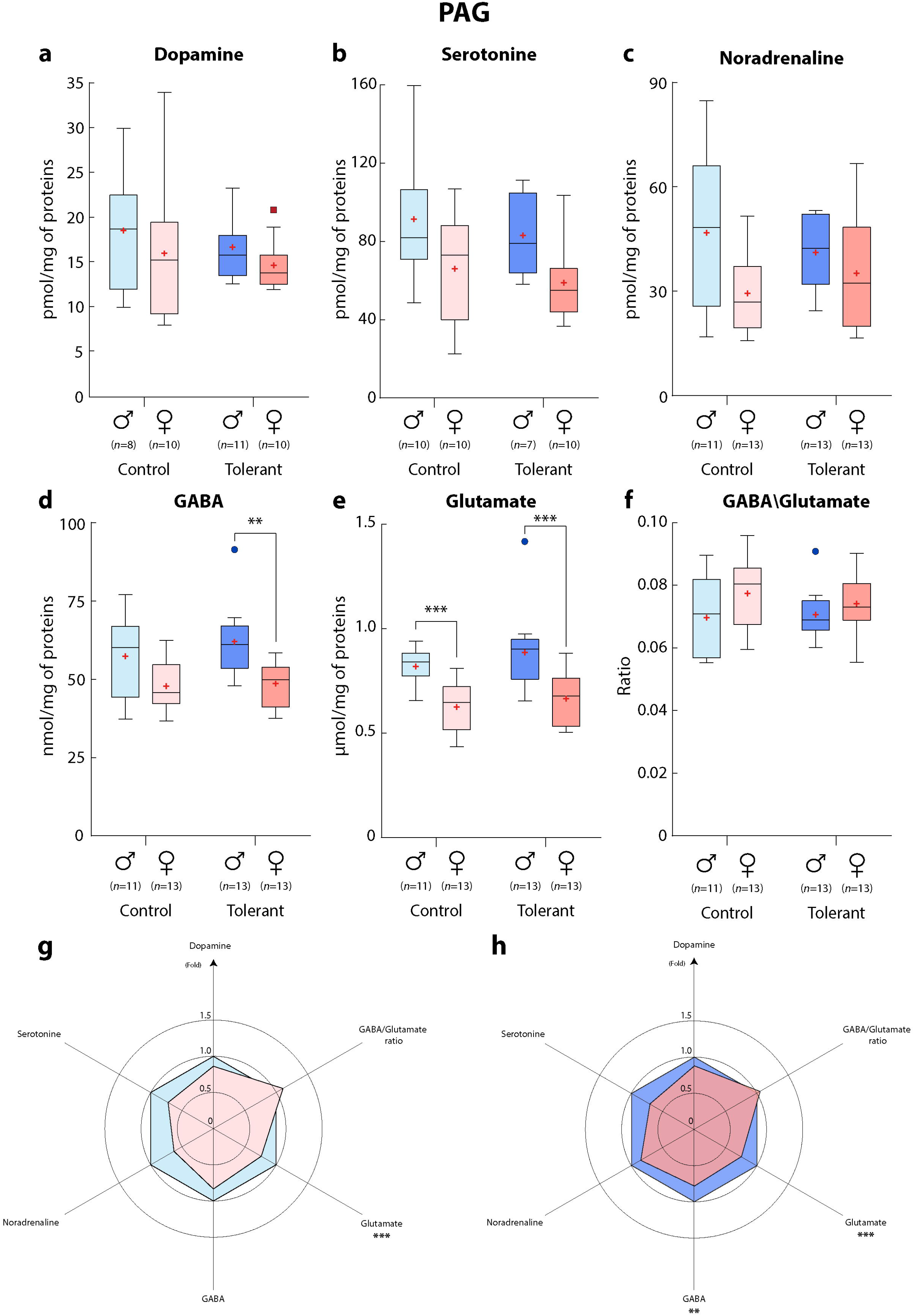
Concentrations of neurotransmitters in the PAG of male and female mice treated with acute or chronic morphine. (**a**) dopamine, (**b**) serotonin, (**c**) noradrenaline, (**d**) GABA, (**e**) glutamate, (**f**) GABA/glutamate ratios. Normalized quantities (according to male levels for each neurotransmitter) of dopamine, serotonin, noradrenaline, GABA, and glutamate, and GABA/glutamate ratio are presented in (**g**) CT mice and (**h**) TOL. The control group (CT) refers to the acute treatment with morphine, while the tolerant group (TOL) designates the chronic treatment leading to antinociceptive tolerance. Data are expressed as means ± SEM, *n* are indicated in the figure. Two-way ANOVA was applied and followed by Tukey’s multiple comparisons test only if F was significant and there was no variance homogeneity. Tukey’s multiple comparisons results are reported as **, *P*< 0.01; ***, *P*<0.001.

**Figure 4.**
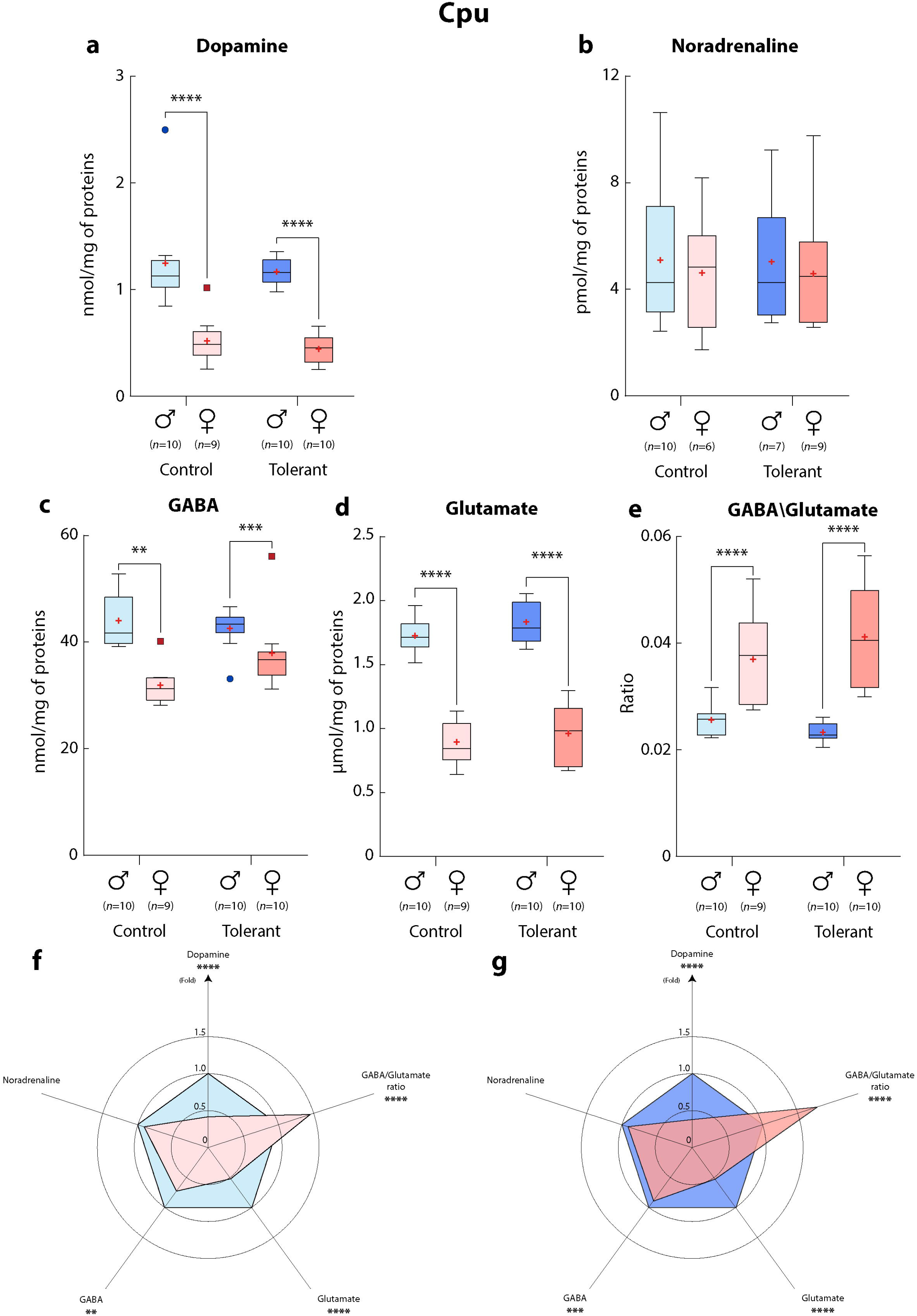
Concentrations of neurotransmitters in the Cpu of male and female mice treated with acute or chronic morphine. (**a**) dopamine, (**b**) noradrenaline, (**c**) GABA, (**d**) glutamate, (**e**) GABA/glutamate ratios. Normalized quantities (according to male levels for each neurotransmitter) of dopamine, serotonin, noradrenaline, GABA, and glutamate, and GABA/glutamate ratio are represented in (**f**) CT mice and (**g**) TOL. The control group (CT) refers to the acute treatment with morphine, while the tolerant group (TOL) designates the chronic treatment leading to antinociceptive tolerance. Data are expressed as means ± SEM, *n* are indicated in the figure. Two-way ANOVA was applied and followed by Tukey’s multiple comparisons test only if F was significant and there was no variance homogeneity. Tukey’s multiple comparisons results are reported as **, *P*<0.01; ***, *P*<0.001; ****, *P*<0.0001.

**Figure 5.**
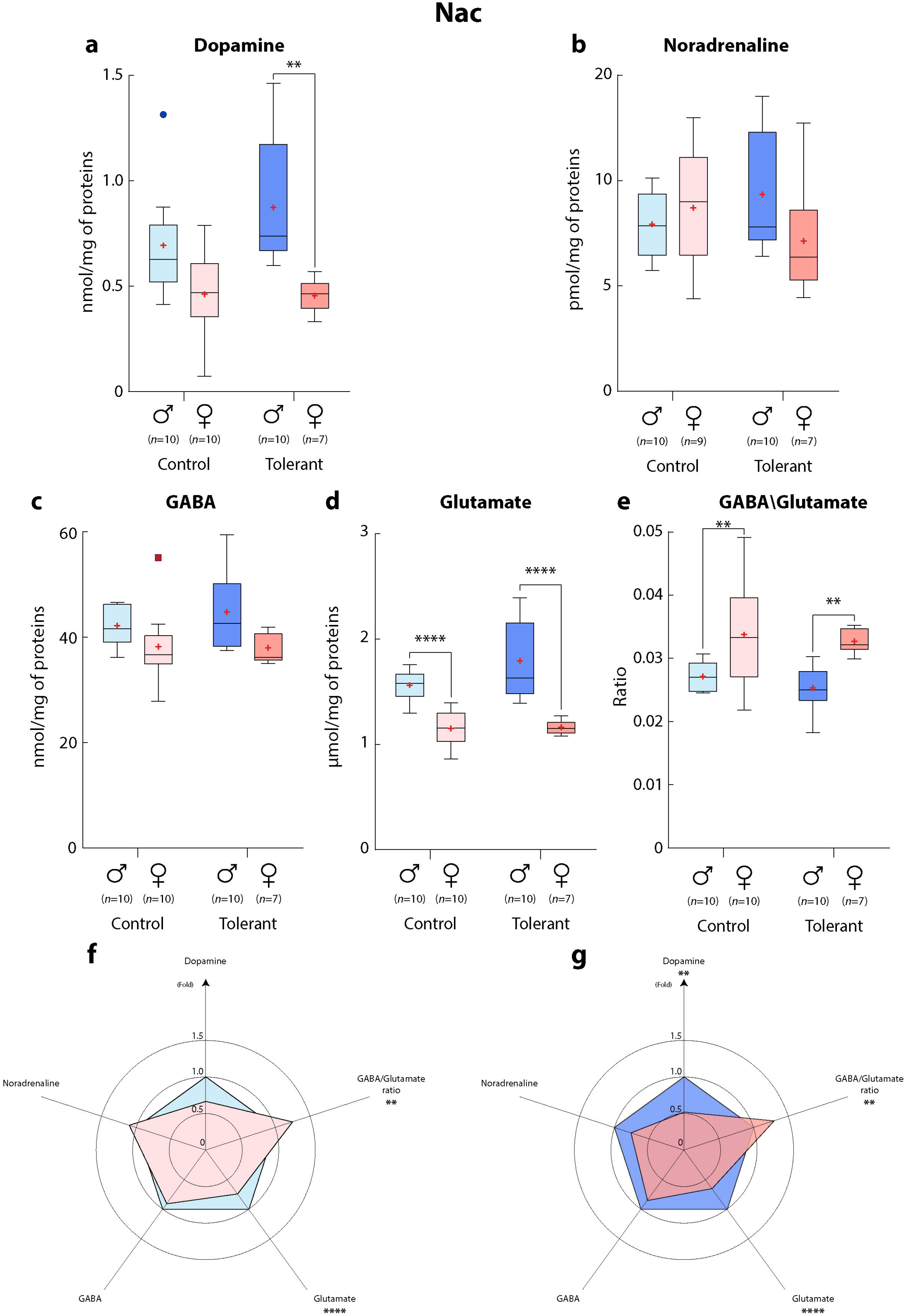
Concentrations of neurotransmitters in the Nac of male and female mice treated with acute or chronic morphine. (**a**) dopamine, (**b**) noradrenaline, (**c**) GABA, (**d**) glutamate, (**e**) GABA/glutamate ratios. Normalized quantities (according to male levels for each neurotransmitter) of dopamine, serotonin, noradrenaline, GABA, and glutamate, and GABA/glutamate ratio in (**f**) CT mice and (**g**) TOL. The control group (CT) refers to the acute treatment with morphine, while the tolerant group (TOL) designates the chronic treatment leading to antinociceptive tolerance. Data are expressed as means ± SEM, *n* are indicated in the figure. Two-way ANOVA was applied and followed by Tukey’s multiple comparisons test only if F was significant and there was no variance homogeneity. Tukey’s multiple comparisons results are reported as **, *P*<0.01; ****, *P*<0.0001.

#### Amygdala

Two-way ANOVA was used to identify the overall effect of sex, treatment and their potential interaction on neurotransmitter amounts in the amygdala (**Figure 2** and **Table S2**). The statistical analysis revealed that significantly higher levels of dopamine (sex, p<0.001) and GABA (sex, p<0.0001) were present in the female amygdala than in the male amygdala, while glutamate levels (sex, p<0.0001) were higher in the male amygdala (**Figure 2a,d,e**). Consequently, the GABA/glutamate ratio (sex, p<0.0001) was significantly higher in females than in males in this brain region (**Figure 2f**). In addition, higher levels of serotonin (treatment, p<0.001) were observed in CT mice compared to TOL mice (**Figure 2b**).

Finally, an interaction between sex and treatment (interaction, p<0.05) was observed in the noradrenaline amounts in the amygdala (**Figure 2c**). Although the post-hoc analysis revealed significantly higher levels of noradrenaline in females than in males in both the CT (Tukey’s multiple comparison test, p<0.001) and TOL mice (Tukey’s multiple comparison test, p<0.0001). Such interaction is probably driven by the effect size which is much higher between CT animals than between TOL animals (**Figure 2c**).

#### Periaqueductal gray

The statistical analysis (**Figure 3** and **Table S2**) revealed that significantly higher levels of serotonin (sex, p<0.01), noradrenaline (sex, p<0.05), GABA (sex, p<0.001) and glutamate (sex, p<0.0001) were present in the male PAG than in the female PAG (**Figure 3b,c,d,e**). No significant difference was observed in the dopamine levels or the GABA/glutamate ratio between the groups (**Figure 3a,f)**. Neither an effect of the treatment nor any interaction was observed in the PAG.

#### Caudate-Putamen

Two-way ANOVA (**Table S2**) showed that significantly higher levels of dopamine and glutamate (sex, p<0.0001) were observed in the male Cpu than in the female Cpu (**Figure 4a,d**). In addition, the GABA/glutamate ratio in the Cpu was found to be significantly higher (sex, p<0.0001) in females than in males (**Figure 4e**). An interaction between sex and treatment (interaction, p<0.05) was observed in the GABA amounts in the Cpu (**Figure 4c**). The post-hoc analysis revealed significantly higher levels of GABA in males than in females in both the CT (Tukey’s multiple comparison test, p<0.01) and TOL (Tukey’s multiple comparison test, p<0.001) groups. As for noradrenaline in the amygdala, the interaction is probably driven by the effect size which is higher between CT animals than between TOL animals (**Figure 4c**).

No effect of the treatment was observed in the Cpu, nor any significant difference in the noradrenaline levels in this brain region (**Figure 4b**). Finally, serotonin levels were below the limit of quantification of our method.

#### Nucleus accumbens

The statistical analysis revealed significant differences between males and females (**Figure 5** and **Table S2)**. Higher levels of dopamine (sex, p<0.0001), GABA (sex, p<0.01) and glutamate (sex, p<0.0001) were observed in the male Nac than in the female Nac (**Figure 5a,c,d**). Contrariwise, the GABA/glutamate ratio was significantly higher in females than in males in this brain region (**Figure 5e**). In addition, higher levels of glutamate (treatment, p<0.05) were observed in CT mice than in TOL mice (**Figure 5d**).

No interaction was observed in the Nac, nor any significant difference in the noradrenaline levels in this brain region (**Figure 5b**). As in the Cpu, serotonin levels were below the limit of quantification of our method in the Nac.

## DISCUSSION

### Sex differences in morphine antinociceptive tolerance

The sex differences in morphine antinociceptive tolerance observed in our experiment were consistent with our last report [31]. As previously seen, although significant differences were observed in the development of the antinociceptive tolerance between male and female mice (*i.e*., females became tolerant to morphine earlier than males during the protocol), the rate at which the tolerance developed did not seem to differ, as witnessed by the absence of difference in the hill slope coefficients. This suggests that the sex disparities observed in our experiment might be related to sex differences in the initial morphine antinociceptive effect at the beginning of the protocol rather than to sex-specific mechanisms involved in the induction of morphine tolerance. This is further corroborated by a study in which the initial sex differences in morphine antinociception were taken into account and no difference was observed in the development of tolerance between males and females [41]. Nevertheless, one should note that sex differences in the hill slope coefficients might have been hidden by the cut-off used in the present experiment (*i.e*., males reached the cut-off for the first days of the protocol, thus, the hill-slope coefficient might have been different if the males were allowed to reach higher latencies).

### Methodology for neurotransmitters quantification in males and females

Our study aimed to quantify neurotransmitters and identify differences between male and female mice after acute and chronic morphine administrations. Considering the importance of sex as a biological variable, especially in pain studies [42], all experiments were conducted according to a 2×2 factorial design to examine the effect of sex and treatment, as well as their potential interaction *via* the application of the 2-way ANOVA, and without increasing the number of animals. The statistical test of the interaction shows whether being male or female changes the effect of chronic morphine administration on neurotransmitter levels. In the absence of interaction, this factorial design allows each factor (*e.g*., sex) to be evaluated over two different conditions (*e.g*., acute and chronic conditions) raising more robust conclusions than individual comparisons [43]. However, even though this strategy can identify relatively small effect sizes with sufficient statistical power, the physiological implication of the statistical result still needs to be elucidated.

In our experiment, brain structures were directly punched (*i.e*., not perfused) 30min after an administration of morphine. Therefore, our quantifications reflect the total amounts of neurotransmitters (mostly production and vesicle storage) after the administration of morphine, in opposition to microdialysis studies focusing on the release of neurotransmitters in much lower quantities. In our study, differences in neurotransmitter contents might thus suggest differences in production rather than secreted amounts. For this reason, it is unlikely that the acute morphine injection in CT mice modulated in about 30min the total quantity of neurotransmitters because modulation of neurotransmitter production involves gene regulation and a complex enzymatic cascade. Nevertheless, whether an acute injection of morphine modifies neurotransmitter production in the brain regions of interest in 30min requires further investigation.

The LC-MS/MS approach used in this study applies the isotopic dilution method for absolute quantification of the neurotransmitters. This approach consists of the addition of a known amount of an internal standard (the molecule of interest bearing one or several rare stable isotopes such as deuterium atoms instead of hydrogen atoms) directly to each sample at the beginning of the sample treatment. Such quantification is based on a signal ratio between the intensity of the molecule of interest and the intensity of the known concentration of the internal standard rather than relying on an external standard curve [34]. This method determines the concentration of the molecule of interest with accuracy, independently of the matrix/tissue context.

In the present article, we used a standard method to produce a comprehensive quantification of most of our targeted neurotransmitters in key brain regions of male and female mice using a robust LC-MS/MS method. In addition, our approach was designed to provide enough statistical power to understand the impact of sex on neurotransmitter levels during acute and chronic treatment with morphine. In the literature, several studies have successfully attempted to quantify neurotransmitters in the same brain regions [44–51]. However, in most cases, the levels of a single neurotransmitter were assessed in a unique brain region of male animals, and/or the methodology differs markedly between studies, yielding difficult comparisons to interpret. For instance, Ongali *et al*., have reported the dopamine levels in the male amygdala using HPLC coupled with electrochemical detection [44]. They have found 7.86 ± 0.97 ng of dopamine/mg of proteins (*i.e*., 51.3 ± 6.4 pmol/mg of proteins) in this brain region, while using our method, we have observed 53.1 ± 6.6 pmole of dopamine/mg and 74.8 ± 7.3 pmole dopamine/mg of protein of dopamine for male and female mice acutely treated with morphine, respectively. Even though these results appear consistent across studies using different methodologies, it is worth noting that Ongalia *et al*., have used B6C3FE-a/a, whereas C57BL/6J mice were used in the present study. These results could suggest no specie difference in the dopamine levels in the amygdala between these mouse strains. However, direct comparisons are hazardous as quantification methodologies are different. Another example is illustrated by comparing our study with the one of Su *et al*.. In this case, Su *et al*., used a similar LC-MS/MS approach than in the present study to quantify most of the neurotransmitters in key brain regions involved in methamphetamine-induced conditioned place preference in C57BL/6J male mice [49]. Noradrenaline levels in the Cpu were reported to be around 0.05 μg/g of wet tissue (*i.e*., 0.30 pmol/mg of wet tissue) in control mice while in our study, we observed 5.09 ± 0.8 and 4.62 ± 0.9 pmol/mg of protein of noradrenaline in the Cpu of male and female CT mice respectively. However, the normalization used in Su *et al*. study and ours tend to yield difficult comparisons.

### Potential origin of sex differences in neurotransmitters levels

Morphine’s impact on neurotransmitter release is widely described in the literature (for review [39]). In addition, morphine administration might lead to modulation of the amounts of intracellular neurotransmitters. The impact of acute and chronic morphine on neurotransmitter total contents in the brain was only sparsely studied. In the present study, we were expecting that, at least, chronic morphine treatment will lead to major changes in neurotransmitter levels in the nociceptive system. However, only a weak effect of the treatment was observed for serotonin (*i.e*., a decrease) in the amygdala. Such surprising results were shaded by the effect of sex on most of the neurotransmitter levels observed through the brain areas investigated in our study.

Sex differences in the production of neurotransmitters might rely on activational and organizational effects of hormones (for review on sex hormones and neurotransmitters, see [52]. For instance, estrogen has been shown to modulate serotonin production through the regulation of the tryptophan hydroxylase [53, 54], the main enzyme responsible for serotonin synthesis. Interestingly, the direction of this effect seems to depend on the brain areas in which estrogen acts [52]. Sex hormones might also have a potent effect on the degradation and uptake of neurotransmitters as well [54–56](Koldzic-Zivanovic, 2004; Smith, 2004; Gundlah, 2002). Taken together, sex hormones may play a role in the observed differences in neurotransmitter levels between males and females in the several brain regions tested.

Although sex differences in neurotransmitter levels were observed in the present study, one should note that many other factors are involved in the behavioral integration of neurotransmission. Such factors might also be subject to sexual dimorphism and include the expression and functionality of neurotransmitter receptors, neuronal circuitry, glial cell activity and neuromodulator expression and production. The set of data presented in this paper is not intended to interpret differences in any behavioral responses between males and females on its own but illustrates the importance of considering sex as a biological variable in neuroscience studies.

## PERSPECTIVES AND SIGNIFICANCE

Our results revealed major sex differences in dopamine, serotonin, noradrenaline, GABA and glutamate levels in the brain regions of interest. Additional studies are required to understand their behavioral meaning but these results illustrate the importance of considering sex as a biological variable in neuroscience studies. Surprisingly, we failed to observe any variations of neurotransmitter concentrations between the acute and chronic morphine conditions, except for serotonin levels in the amygdala which slightly decreased during chronic treatment with morphine. These results suggest a limited impact of morphine chronic treatment on neurotransmitter production within the brain regions of interest.

## ABBREVIATIONS

5-HT_2C_ receptor: 5-hydroxy tryptamine receptor_2C_
ACN: acetonitrile
CID: collision gas
CNS: central nervous system
Cpu: caudate-putamen
CT: group of mice treated acutely with morphine
ED50: half-maximal effective dose
GABA: γ-Aminobutyric acid
IS: internal standard
LC-MS/MS: liquid chromatography coupled to tandem mass spectrometry
LSC: lumbar spinal cord
MOR: mu-opioid receptor
MPE: maximal possible effect
MRM: multiple reaction monitoring mode
Nac: nucleus accumbens
PAG: periaqueductal gray
TIT: tail immersion test
TOL: group of mice groups treated chronically with morphine
UGT: UDP-glucuronosyl-transferase
VTA: ventral tegmental area

## ACKNOWLEDGMENTS AND GRANT SPONSOR

This work was funded by CNRS, University of Strasbourg (Unistra) and French Ministère Délégué à la Recherche et à l’Enseignement Supérieur (PhD fellowship to F.G., A.-K.B. and V.H.). We thank the following research programs of excellence for their support: FHU Neurogenycs, French National Research Agency (ANR) through the Programme d’Investissement d’Avenir (contract ANR-17-EURE-0022, EURIDOL graduate school of pain).

## AUTHOR CONTRIBUTIONS

**Conceptualisation**, Y.G., F.G., V.H.; **Methodology**, Y.G., F.G., V.H., V.A.; **Investigation**, F.G., V.H., V.A. A.-K.B.; **Writing – Original Draft**, Y.G., F.G.; **Writing – Review & Editing**, Y.G., F.G.; **Funding Acquisition**, Y.G.; **Resources**, Y.G.; **Supervision**, Y.G.

## DATA AVAILABILITY

The data that support the findings of this study are available from the corresponding author upon request.

## CONSENT FOR PUBLICATION

All authors have agreed to publish this manuscript

## DECLARATION OF TRANSPARENCY AND SCIENTIFIC RIGOUR

This Declaration acknowledges that this paper adheres to the principles for transparent reporting and scientific rigour of preclinical research as stated in the BJP guidelines for Design and Analysis, and Animal Experimentation, and as recommended by funding agencies, publishers and other organisations engaged with supporting research.

## LEGENDS

### TABLES

**Table S1. LC-MS/MS conditions.** LC and MS/MS conditions for the purification, detection and quantification of morphine and M3G and their respective heavy-tagged counterparts. The flow rate was set at 90 μl/min on a ZORBAX SB-C18 column (150 x 1mm, 3.5μm).

**Table S2. Statistical details for morphine antinociceptive effect and induction of tolerance (see Figure 1).** Non-linear regression with a 4-parameters logistic equation was applied to define the ED_50_ of morphine and the 95% confidence intervals in both males and females. The two fits were compared using a nested-model comparison with the extra sum-of-the-squares F test.

For the tolerance experiment, the same analysis was applied to the data of each mouse. Then, the obtained parameters were averaged and compared with an unpaired t-test with Welch’s correction. MPE, maximal possible effect.

**Table S3. Statistical details for concentrations of dopamine, serotonin, noradrenaline, GABA and glutamate in the amygdala, PAG, Cpu and Nac of male and female mice treated with acute or chronic morphine (see Figure 2–5).** Two-way ANOVA was used to identify the differences between each group.

