## Supplementary Tables files for "Sex differences in neurotransmitter levels in different brain regions after acute and chronic morphine treatment in mice"

Abbreviated title: Neurotransmiters levels are driven by sex.

**Florian Gabel^1^, Volodya Hovhannisyan^1^, Abdel-Karim Berkati^1^,**

**Virginie Andry^1,2^ & Yannick Goumon^1,2*^**

***Tables***

**Table S1. LC-MS/MS conditions.** LC and MS/MS conditions for the purification, detection and quantification of dopamine, serotonin, noradrenaline, GABA and glutamate, as well as their respective heavy-tagged counterparts. The flow rate was set at 90 µl/min on a ZORBAX SB-C18 column (150 x 1mm, 3.5μm).

**Table S2. Statistical details for morphine antinociceptive effect and induction of tolerance (see Figure 1).** Non-linear regression with a 4-parameters logistic equation was applied to define the ED_50_ of morphine and the 95% confidence intervals in both males and females. The two fits were compared using a nested-model comparison with the extra sum-of-the-squares F test.

For the tolerance experiment, the same analysis was applied to the data of each mouse. Then, the obtained parameters were averaged and compared with an unpaired t-test with Welch’s correction. MPE, maximal possible effect.

**Table S3. Statistical details for the quantification of dopamine, serotonin, noradrenaline, GABA and glutamate in the amygdala, PAG, caudate-putamen, nucleus accumbens and lumbar spinal cord.** Two-way ANOVA was used to assess the differences in dopamine, serotonin, noradrenaline, GABA and glutamate quantities between the groups.

**Table S1**.

|  | **Mobile phase** | | |
| --- | --- | --- | --- |
|  | **ACN** | **H_2_O** | **Formic acid** |
| **Mobile phase A** | 1% | 98.9% | 0.1% |
| **Mobile phase B** | 99.9% | 0 | 0.1% |

| **HPLC gradient** | | | | | | |  |
| --- | --- | --- | --- | --- | --- | --- | --- |
| **Time (min)** | 0 | 2.5 | 9 | 11 | 12 | 12.5 | 18 |
| **% B mobile phase** | 0 | 0 | 20 | 98 | 98 | 0 | 0 |

| **MS parameters** | |
| --- | --- |
| **Mode** | positive |
| **Spray voltage** | 3,500 V |
| **Nebulizer gas** | Nitrogen |
| **Desolvation (nitrogen) sheath gas** | 18 Arb |
| **Aux gas** | 7 Arb |
| **Ion transfer tube temperature** | 297°C |
| **Vaporizer temperature** | 131°C |
| **Q1 and Q3 resolutions** | 0.7 FWHM |
| **Collision gas (CID, argon) pressure** | 2 mTorr |

**MS ionization, selection, fragmentation and identification parameters**

| **Compound** | **Polarity** | **Precursor**  **(m/z)** | **Product**  **(m/z)** | **Ion product**  **type** | **Collision Energy (V)** | **RF Lens (V)** |
| --- | --- | --- | --- | --- | --- | --- |
| **Dopamine** | Positive | 324.14 | 116.11 | Quantification | 54.24 | 206 |
| **Dopamine** | Positive | 324.14 | 145.11 | Qualification | 33.36 | 206 |
| **Dopamine** | Positive | 324.14 | 171.06 | Qualification | 23.90 | 206 |
| **D4-Dopamine** | Positive | 328.14 | 122.11 | Quantification | 35.00 | 206 |
| **Noradrenaline** | Positive | 340.15 | 116.11 | Quantification | 55.00 | 215 |
| **Noradrenaline** | Positive | 340.15 | 145.11 | Qualification | 36.19 | 215 |
| **Noradrenaline** | Positive | 340.15 | 171.06 | Qualification | 24.61 | 215 |
| **C6-Noradrenaline** | Positive | 346.15 | 145.11 | Quantification | 36.04 | 226 |
| **C6-Noradrenaline** | Positive | 346.15 | 158.11 | Qualification | 18.34 | 226 |
| **C6-Noradrenaline** | Positive | 346.15 | 171.06 | Qualification | 25.62 | 226 |
| **Serotonine** | Positive | 347.13 | 145.06 | Quantification | 18.49 | 232 |
| **Serotonine** | Positive | 347.13 | 160.07 | Qualification | 20.82 | 232 |
| **Serotonine** | Positive | 347.13 | 171.00 | Qualification | 27.54 | 232 |
| **D4-Serotonine** | Positive | 351.16 | 145.04 | Quantification | 19.51 | 219 |
| **D4-Serotonine** | Positive | 351.16 | 164.06 | Qualification | 21.38 | 219 |
| **D4-Serotonine** | Positive | 351.16 | 351.16 | Qualification | 28.10 | 219 |
| **GABA** | Positive | 274.11 | 116.04 | Quantification | 46.35 | 157 |
| **GABA** | Positive | 274.11 | 128.11 | Qualification | 43.47 | 157 |
| **GABA** | Positive | 274.11 | 171.06 | Qualification | 19.66 | 157 |
| **D6-GABA** | Positive | 280.16 | 116.11 | Quantification | 48.73 | 150 |
| **D6-GABA** | Positive | 280.16 | 128.04 | Qualification | 44.78 | 232 |
| **D6-GABA** | Positive | 280.16 | 171 | Qualification | 19.60 | 232 |
| **Glutamate** | Positive | 318.11 | 116.11 | Quantification | 55.00 | 169 |
| **Glutamate** | Positive | 318.11 | 128.11 | Qualification | 52.82 | 169 |
| **Glutamate** | Positive | 318.11 | 170.93 | Qualification | 20.52 | 169 |
| **D5-Glutamate** | Positive | 323.12 | 116.11 | Quantification | 55 | 159 |
| **D5-Glutamate** | Positive | 323.12 | 128.06 | Qualification | 48.68 | 159 |
| **D5-Glutamate** | Positive | 323.12 | 171.00 | Qualification | 20.77 | 159 |

**Table S2**

|  | **Behavioural experiments** | |
| --- | --- | --- |
|  | **Tolerance experiment** | |
| Parameters | **Time at which half %MPE** | **Hill-slope coefficient** |
| Test | Unpaired t-test | Unpaired t-test |
| F, t, df | t=15.06 df=10 | t=0.9031 df=10 |
| p-value | < 0.0001* | 0.3877 |

**Table S3.**

|  | **CNS regions quantification** | | | | | |
| --- | --- | --- | --- | --- | --- | --- |
|  | **Amygdala** | | | | | |
|  | **Dopamine** | **Serotonin** | **Noradrenaline** | **GABA** | **Glutamate** | **Ratio GABA/**  **glutamate** |
| Interaction | F (1, 48) = 0.01551  P=0.9014 | F (1, 45) = 1.393  P=0.2441 | F (1, 48) = 6.756  P=0.0124* | F (1, 48) = 1.938  P=0.1703 | F (1, 48) = 0.05356  P=0.8180 | F (1, 48) = 0.5643  P=0.4562 |
| Treatment | F (1, 48) = 2.280  P=0.1376 | F (1, 45) = 12.68  P=0.0009* | F (1, 48) = 0.2141  P=0.6447 | F (1, 48) = 0.1485  P=0.7017 | F (1, 48) = 0.6816  P=0.4131 | F (1, 48) = 1.422  P=0.2389 |
| Sex | F (1, 48) = 13.67  P=0.0006* | F (1, 45) = 2.969  P<0.0917 | F (1, 48) = 41.51  P<0.0001* | F (1, 48) = 59.34  P<0.0001* | F (1, 48) = 23.37  P<0.0001* | F (1, 48) = 111.1  P<0.0001* |
|  | **PAG** | | | | | |
|  | **Dopamine** | **Serotonin** | **Noradrenaline** | **GABA** | **Glutamate** | **Ratio GABA/**  **glutamate** |
| Interaction | F (1, 35) = 0.1402  P=0.7104 | F (1, 33) = 0.003638  P=0.9523 | F (1, 46) = 0.5851  P= 0.4482 | F (1, 46) = 0.4986  P=0.4837 | F (1, 46) = 0.01947  P=0.8896 | F (1, 46) = 0.5313  P=0.4697 |
| Treatment | F (1, 35) = 0.4538  P=0.5050 | F (1, 33) = 0.7402  P=0.3958 | F (1, 46) = 0.1388  P=0.7112 | F (1, 46) = 0.9542  P=0.3338 | F (1, 46) = 1.916  P=0.1730 | F (1, 46) = 0.1501  P=0.7003 |
| Sex | F (1, 35) = 3.571  P=0.0671 | F (1, 33) = 7.555  P=0.0096* | F (1, 46) = 6.466  P=0.0114* | F (1, 46) = 16.89  P=0.0002* | F (1, 46) = 37.50  P<0.0001* | F (1, 46) = 3.649  P=0.0624 |
|  | **Caudate-putamen** | | | | | |
|  | **Dopamine** | **Serotonin** | **Noradrenaline** | **GABA** | **Glutamate** | **Ratio GABA/**  **glutamate** |
| Interaction | F (1, 35) = 0.3890  P=0.5369 | / | F (1, 28) = 0.006672  P=0.9355 | F (1, 35) = 4.973  P=0.0323* | F (1, 35) = 0.1318  P=0.7187 | F (1, 35) = 3.763  P=0.0605 |
| Treatment | F (1, 35) = 0.3739  P=0.5449 | / | F (1, 28) = 0.02651  P=0.8718 | F (1, 35) = 2.159  P=0.1507 | F (1, 35) = 2.211  P=0.1460 | F (1, 35) = 0.7519  P=0.3918 |
| Sex | F (1, 35) = 102.5  P<0.0001* | / | F (1, 28) = 0.2228  P<0.6406 | F (1, 35) = 31.42  P<0.0001* | F (1, 35) = 214.7  P<0.0001* | F (1, 35) = 86.03  P<0.0001* |
|  | **Nuleus accumbens** | | | | | |
|  | **Dopamine** | **Serotonin** | **Noradrenaline** | **GABA** | **Glutamate** | **Ratio GABA/**  **glutamate** |
| Interaction | F (1, 33) = 1.377  P=0.2490 | **/** | F (1, 32) = 3.031  P=0.0913 | F (1, 33) = 0.4859  P=0.4906 | F (1, 33) = 2.235  P=0.1444 | F (1, 33) = 0.05578  P=0.8147 |
| Treatment | F (1, 33) = 1.176  P=0.2861 | **/** | F (1, 32) = 0.009221  P=0.9241 | F (1, 33) = 0.3737  P=0.5452 | F (1, 33) = 2.599  P<0.1164 | F (1, 33) = 0.8263  P=0.3699 |
| Sex | F (1, 33) = 16.64  P=0.0003* | **/** | F (1, 32) = 0.6980  P=0.4096 | F (1, 33) = 7.623  P=0.0093* | F (1, 33) = 48.72  P<0.0001* | F (1, 33) = 20,22  P<0.0001* |
